# Chemotherapy drugs induce different gut microbiota disorder pattern and NODs/RIP2/NF-κB signaling pathway activation that lead to different degrees of intestinal injury

**DOI:** 10.1101/2022.05.04.490707

**Authors:** Bin Huang, Mengxuan Gui, Jinyan Zhao, Zhuona Ni, Yanbin He, Jun Peng, Jiumao Lin

**Author notes:** Contributed equally (Bin Huang, Mengxuan Gui). Corresponding author: **Jun Peng (PhD)**, Fujian Key Laboratory of Integrative Medicine on Geriatrics, Fujian University of Traditional Chinese Medicine, Fuzhou, Fujian 350122, P.R. China, E-mail address, **Jiumao Lin (PhD)**, Academy of Integrative Medicine of Fujian University of Traditional Chinese Medicine, 1 Qiuyang Road, Minhou Shangjie, Fuzhou, Fujian 350122, P.R. China.

## Abstract

5-Fluorouracil (5-FU), irinotecan (CPT-11), oxaliplatin (L-OHP) and calcium folinate (CF) are the widely used chemotherapy drugs to treat colorectal cancer. However, the use of chemotherapy is often accompanied by intestinal inflammation and gut microbiota disorder. Moreover, the change of gut microbiota may lead to destruction of the intestinal barrier, which contributes to the severity of intestinal injury. There was no detailed comparison of intestinal injury and gut microbiota disorder among 5-FU, CPT-11, L-OHP and CF, which is not benefit for the development of targeted detoxification therapy after chemotherapy. In this project, a model of chemotherapy-induced intestinal injury in tumor-bearing mice was established by intraperitoneal injection of chemotherapy drugs at a clinically equivalent dose. 16S rDNA sequencing was used to detect gut microbiota. We found that 5-FU, CPT-11 and L-OHP caused intestinal injury, inflammatory cytokine (IFN-γ, TNF-α, IL-1β, and IL-6) secretion, and gut microbiota disorder. Importantly, we established a complex but clear network between the gut microbiota change pattern and intestinal damage degree induced by different chemotherapy drugs. L-OHP caused the most severe damage in intestine and disorder of gut microbiota, and showed considerable overlap of the microbiota change pattern with 5-FU and CPT-11. The phylogenetic investigation of communities by reconstruction of unobserved states, V1.0 (PICRUSt) analysis showed that the microbiota disorder pattern induced by 5-FU, CPT-11 and L-OHP was related to the NOD like signaling pathway. Therefore, we detected the protein expression of the NODs/RIP2/NF-κB signaling pathway and found that L-OHP activated that pathway highest. Furthermore, by RDA/CCA analysis, we found that *Bifidobacterium, Akkermansia, Allobaculum, Catenibacterium, Mucispirillum, Turicibacter*, *Helicobacter, Proteus, Escherichia Shigella, Alloprevotealla, Vagococcus, Streptococcus* and *Candidatus Saccharimonas* were highly correlated with the NODs/RIP2/NF-κB signaling pathway, and influenced by chemotherapy drugs.

**IMPORTANCE:** The chemotherapy-induced intestinal injury limit drugs clinical use. Intestinal injury involves multiple signaling pathways and the disruption of microbiota. Our results suggest that the degree of intestinal injury caused by different drugs of the first-line colorectal chemotherapy regimen is related to the change pattern of microbiota. Moreover, the NODs/RIP2/NF-κB signaling pathway was activated in different degrees is also related to the change pattern of microbiota. We found L-OHP caused the most severe change of gut microbiota, and showed considerable overlap of the microbiota changes pattern with 5-FU and CPT-11. Here, we have established a network of different chemotherapy drugs, gut microbiota and NODs/RIP2/NF-κB signaling pathway, which may provide a new basis for further elucidating the mechanism and clinical treatment of intestinal injury caused by chemotherapy.

---

**C**ancer is a major public health problem. Colorectal cancer is the third highest incidence and the fifth highest mortality rate among cancer (1). Few patients with colorectal cancer are diagnosed at clinical stage I, most are diagnosed in the advanced stages (2, 3). It is difficult to eliminate colorectal cancer by surgical resection, only because of the high probability of tumor recurrence and metastasis. Due to the development of neoadjuvant and adjuvant chemotherapy, patients can take appropriate chemotherapy drugs for adjuvant treatment before or after surgery to improve the surgical resection rate and prevent tumor recurrence or metastasis (3). 5-Fluorouracil (5-FU), irinotecan (CPT-11), oxaliplatin (L-OHP) and calcium folinate (CF) are the most widely used chemotherapy drugs to treat colorectal cancer (4). 5-FU, CPT-11 and L-OHP inhibit tumor cell proliferation and induce apoptosis by interfering with DNA and RNA synthesis in tumor cells (5). CF has no antitumor activity, but is commonly used in FOLFOX to increase the efficacy of 5-FU (6).

The interference of chemotherapy in DNA and RNA synthesis targets not only tumor cells but also normal cells, which leads to a series of side effects. Approximately 50-80% of patients with colorectal cancer treated with 5-FU, CPT-11, or L-OHP develop chemotherapy-induced intestinal injury with severe diarrhea, nausea, vomiting, anorexia, and weight loss (7). The debilitating effects of intestinal injury include pain, increased length of hospitalization, decreased quality of patient life, modification of anti-neoplastic treatment regimens, higher risk of systemic infections, and even death (8-10).

Several contributing pathogenic elements of chemotherapy-induced intestinal injury have been identified, including crypt epithelium apoptosis, hypoproliferation, abnormal inflammation (11, 12) and gut microbiota disorder (13, 14). Changes in the gut microbiota may cause destruction of the intestinal barrier, then cause serious intestinal injury (15). The Nucleotide Binding Oligomerization Domain Containing 1(NOD1) and NOD2 can be activated by exogenous microbiota, as well as the NODs/receptor-interacting protein (RIP2)/nuclear factor (NF)-κB signaling pathway plays an essential role in inflammation (16). After activation of this pathway, the secretion of several inflammatory cytokines, including tumor necrosis factor (TNF)-α, interleukin-1β (IL-1β), IL-6, and interferon-γ (IFN-γ), are up-regulated (17). Therefore, the aim of this study was to investigate whether chemotherapy drugs (5-FU, CPT-11, L-OHP, and CF) would lead to different pathophysiology of intestinal injury, patterns of gut microbiota disorder, and the molecular mechanisms. Understanding these can provide new ideas and directions for the targeted treatment of colorectal cancer chemotherapy-induced intestinal injury.

## RESULTS

### The influence of different chemotherapy drugs on mice mortality, body weight loss, diarrhea index, and fecal occult blood (FOB) score

Model established and the mortality of the mice was observed and recorded (Fig. 1a). In the L-OHP group, mice died from Day 3 onward, with two mice dying overall. In the 5-FU or CPT-11 group, only one mouse died on Day 2, respectively (Fig. 1b). Meanwhile, all mice remained alive on Day 5 in CF and control groups. 5-FU, CPT-11 and L-OHP effectively inhibited tumor growth (Fig. 1c). While, contrary to the slight increase in body weight in control and CF groups, body weight loss was observed in 5-FU, CPT-11 and L-OHP groups (Fig. 1d). The weights of mice in control and CF groups increased by 6.66% and 5.68% (Day 5 vs. Day 1), respectively. However, the weights decreased by 15.05%, 17.61%, and 27.55% in the 5-FU, CPT-11, and L-OHP groups, respectively. After treatment with 5-FU, CPT-11 or L-OHP for 5 days, the diarrhea index, FOB score, and disease activity index (DAI) were substantially higher compared to those of the control group. The mean diarrhea index reached 1.86, 1.71 and 2.33 (Fig. 1e); the mean FOB scores reached 2.00, 2.00, and 2.67 (Fig. 1f); and the mean DAI reached 4.86, 5.14 and 7.17 (Fig. 1g) for 5-FU, CPT-11 and L-OHP, respectively. Moreover, no significant differences of diarrhea index, DAI and FOB were observed between the CF and control groups.

**FIG 1.**
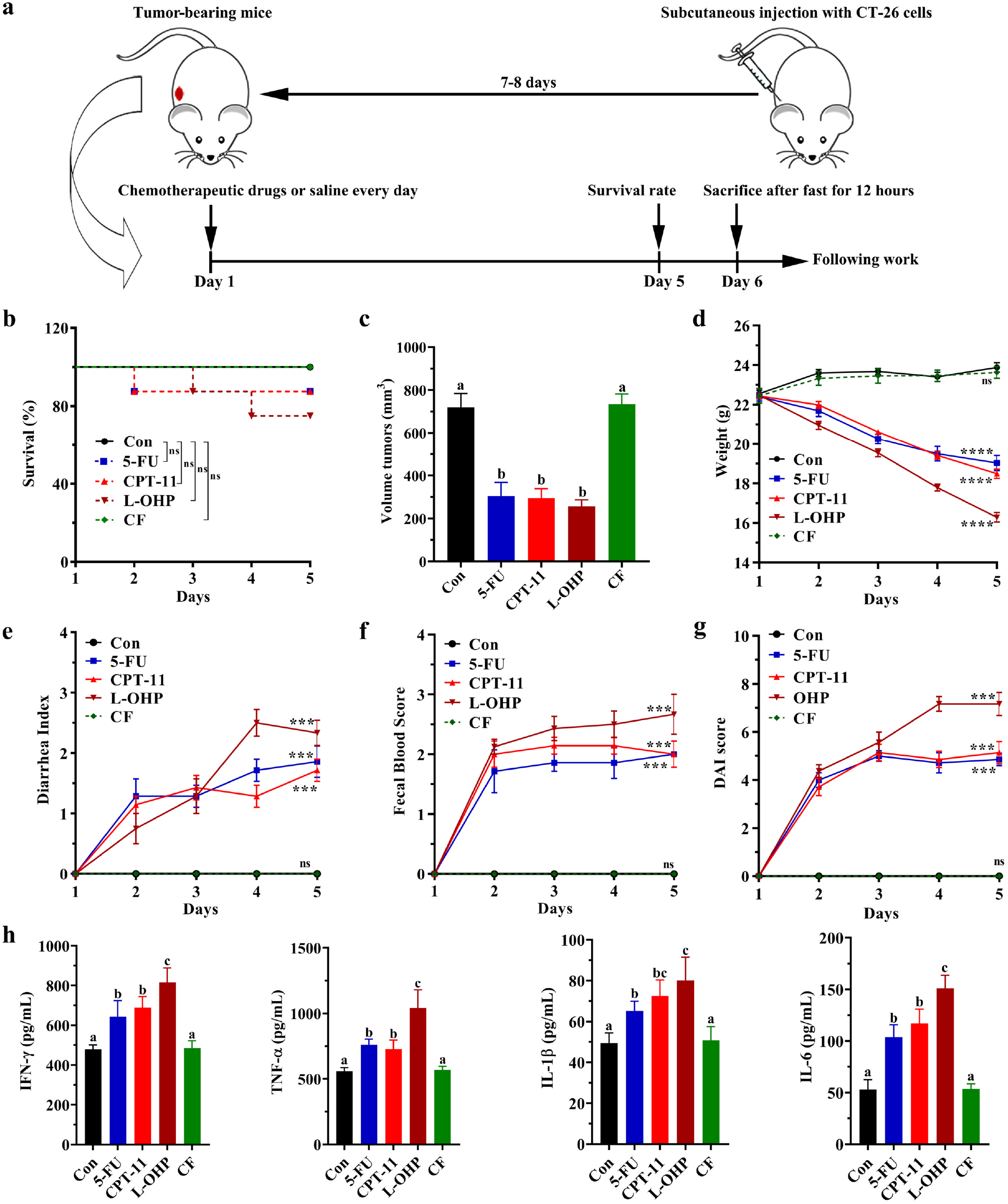
Different chemotherapy drugs induced intestinal mucositis in mice. (a) Experimental procedure for intestinal mucositis model administration. (b to d) The mortality (b), tumor volume (c) and body weight (d) of the mice indifferent groups. (e to g) The average diarrhea index (e), fecal blood score (f) and DAI (g) of the mice. (h) The expression of inflammatory cytokines (IFN-γ, TNF-α, IL-1β, and IL-6) in the serum of mice. The data present the mean ± standard deviation (SD) (n = 8). One-way ANOVA followed by Tukey’s test was used to evaluate the statistical significance. The different letters represent significant differences between different groups (p < 0.05).

### The secretion of inflammatory cytokines in the serum of mice treated with different chemotherapy drugs

5-FU, CPT-11, and L-OHP administration significantly enhanced the secretion of inflammatory cytokines, including IFN-γ, TNF-α, IL-1β, and IL-6 in the serum of mice compared to the control group (p < 0.05) (Fi. 1h). The levels of cytokines were not significantly different between the CF and control groups (p > 0.05).

### The damage of different chemotherapy drugs to the liver, spleen, kidneys and intestines

We investigated the toxicological damage of different chemotherapy drugs to the liver, spleen and kidneys. 5-FU, CPT-11 and L-OHP reduced the weight of the liver, spleen and kidneys. The weight decreased more in the L-OHP group than in the 5-FU or CPT-11 groups (p < 0.05) (Fig. 2a to c). While, only 5-FU and L-OHP reduced the liver and spleen index (tissues index = weight of tissues/weight of body) compared to the control group (p < 0.05) (Fig. 2a to c). Only the L-OHP group showed significantly histopathological damage in the liver and spleen (Fig. S1).

**FIG 2.**
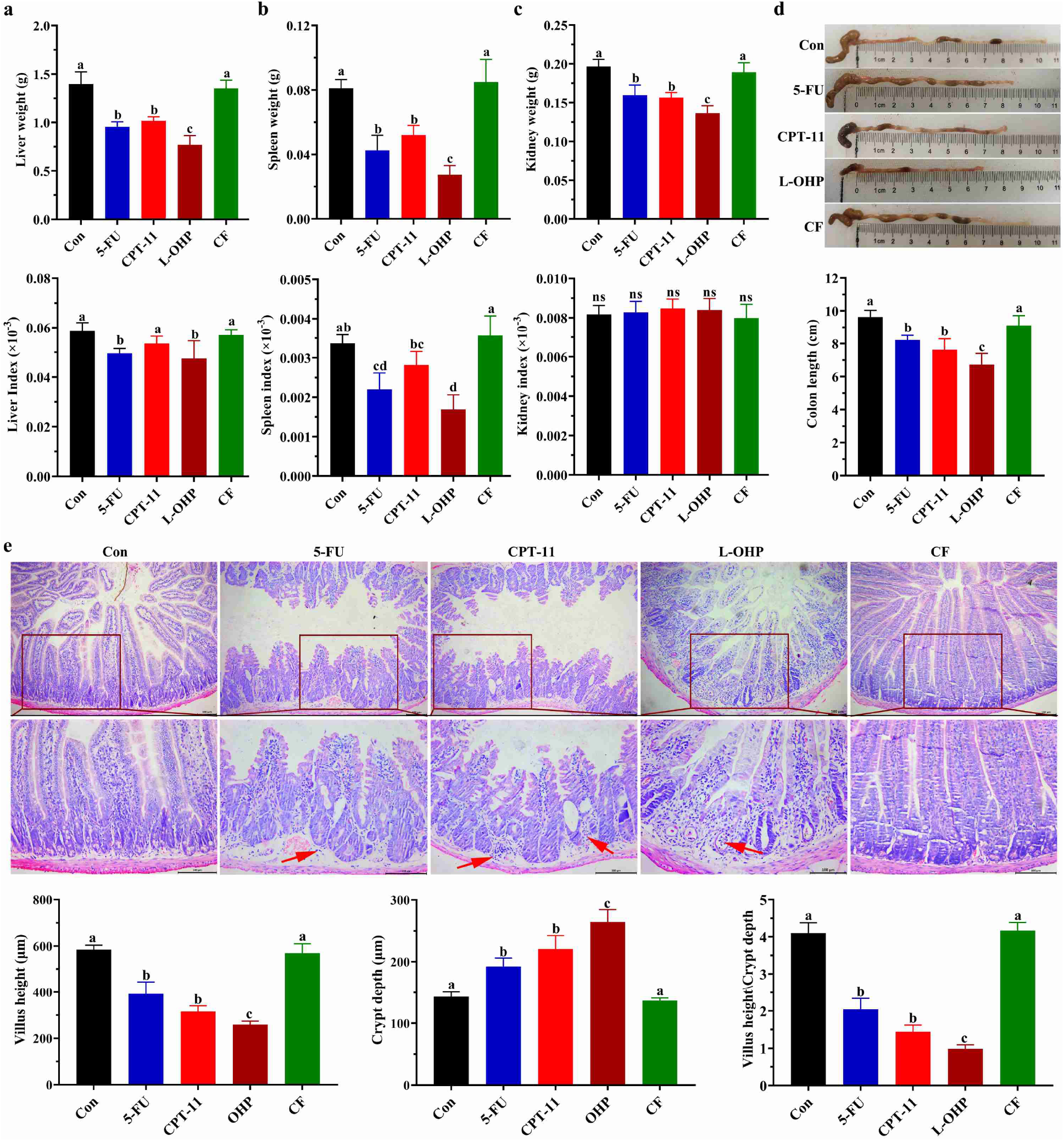
Different chemotherapy drugs caused organ leison in mice. (a) Graphs of average liver weight and liver index in the different groups on Day5. (b) Graphs of average spleen weight and spleen index of the different groups on Day 5. (c) Graphs of average kidney weight and kidney index in the groups on Day 5. (d) Graphs of average colon length and a photograph for measuring and comparing colon length in the groups on Day 5. (e) Hematoxylin and Eosin (HE) staining of representative histological sections of the jejunum from the groups (200× magnification), and the villus height, crypt depth and villus/crypt ratio were measured. One-way ANOVA followed by Tukey’s test was used to evaluate the statistical significance. The different letters represent significant differences between different groups (p < 0.05).

The intestinal injury induced by chemotherapy drugs was the focus of this study. We found that L-OHP, CPT-11 and 5-FU shortened the length of the colon, but not CF (Fig. 2d). The results of histopathology tests showed that the intestines of the mice in the control and CF groups maintained an intact structure, while obvious histological changes, such as shortening of the villi and the hyperplasia of crypts of jejunum (Fig. 2e), as well as inflammatory cell infiltration and crypt large area loss in the colon (Fig. S2) were observed in 5-FU, CPT-11 and L-OHP groups. Moreover, the histopathology results showed that the L-OHP group induced the most serious intestine damage than 5-FU and CPT-11 groups.

### Cellular proliferation and apoptosis of jejunum under different chemotherapy drugs treatment

Chemotherapy drugs can induce damage to the jejunum and colon, especially in the jejunum (18, 19). Therefore, we used the jejunum as the main tissue for the subsequent experiment. PCNA expression, an index of cell proliferation, was detected. The number of PCNA^+^ cells significantly decreased in the 5-FU, CPT-11 and L-OHP groups (vs. control, p < 0.05), and the decrease was dramatic in the L-OHP group (vs. 5-FU and vs CPT-11, p < 0.05) (Fig. 3a). However, there was no significant decrease in the CF group compared to the control group (p < 0.05). As expected, the results of TUNEL staining were contrary to PCNA staining (Fig. 3b).

**FIG 3.**
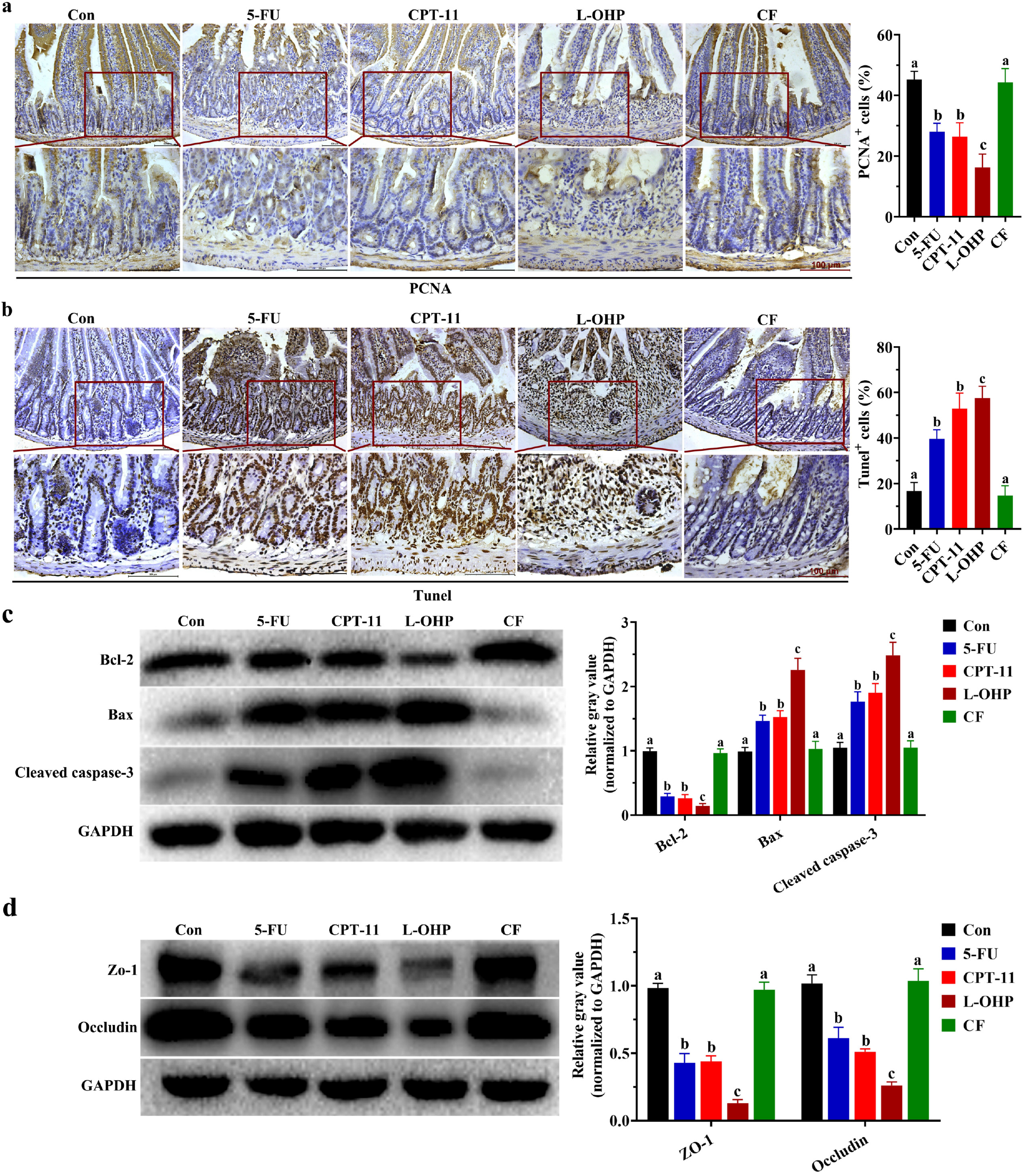
Effect of different chemotherapy drugs on apoptosis and proliferation. (a) Immunohistochemical (IHC) staining was used to detect the expression of PCNA and the average PCNA-positive cells proportions. (b) TUNEL assay for determining the DNA damage and the average apoptosis-positive cell proportions. (c and d) The protein expressions of cleaved-caspase 3, Bcl-2, Bax (c), and Zo-1, Occludin (d). The protein production was normalized with GAPDH. One-way ANOVA followed by Tukey’s test was used to evaluate the statistical significance. The different letters represent significant differences between different groups (p < 0.05).

To further study the apoptotic response in the jejunum, the levels of apoptosis-related molecules, such as Bcl-2, Bax and cleaved caspase-3 were detected. As shown in Fig. 3c, administration of 5-FU, CPT-11 and L-OHP significantly reduced the protein level of Bcl-2, but increased the protein levels of Bax and cleaved caspase-3 (p < 0.05). Noticeably, the change of these proteins were more obviously in the L-OHP group than in the 5-FU or CPT-11 groups (p < 0.05). The protein levels of these molecules were not significantly different between the CF and control groups (p > 0.05).

### The Expression of the jejunal mucosal barrier protein ZO-1 and occludin in mice treated with different chemotherapy drugs

We examined the expression of ZO-1 and occludin, which are associated with the integrity and permeability of the jejunal mucosal barrier (Fig. 3d). The protein levels of ZO-1 and occludin were markedly decreased in 5-FU, CPT-11 and L-OHP groups, and the decrease was more significantly in the L-OHP group than 5-FU or CPT-11 (p < 0.05). However, treatment with CF resulted in no significant changes in ZO-1 and occludin expression compared to the control group (p > 0.05).

### Changes in gut microbiota under different chemotherapy drug treatment

Intestinal microbiology serves as an important regulator of intestinal health. Disturbance of gut microbiota can promote intestinal injury (20). Intestinal injury caused by chemotherapy drugs can be effectively alleviated by adjusting the gut microbiota (21). However, no comparison of the gut microbiota disorder pattern has been made among different colorectal cancer chemotherapy drug, which makes it difficult to treat intestinal injury induced by chemotherapy. In this study, we performed a systematic sequencing analysis by detecting the 16S rRNA gene in variable regions V3-V4 to study the alteration of the gut microbiota from fecal samples in all groups (Fig. 4a to e).

**FIG 4.**
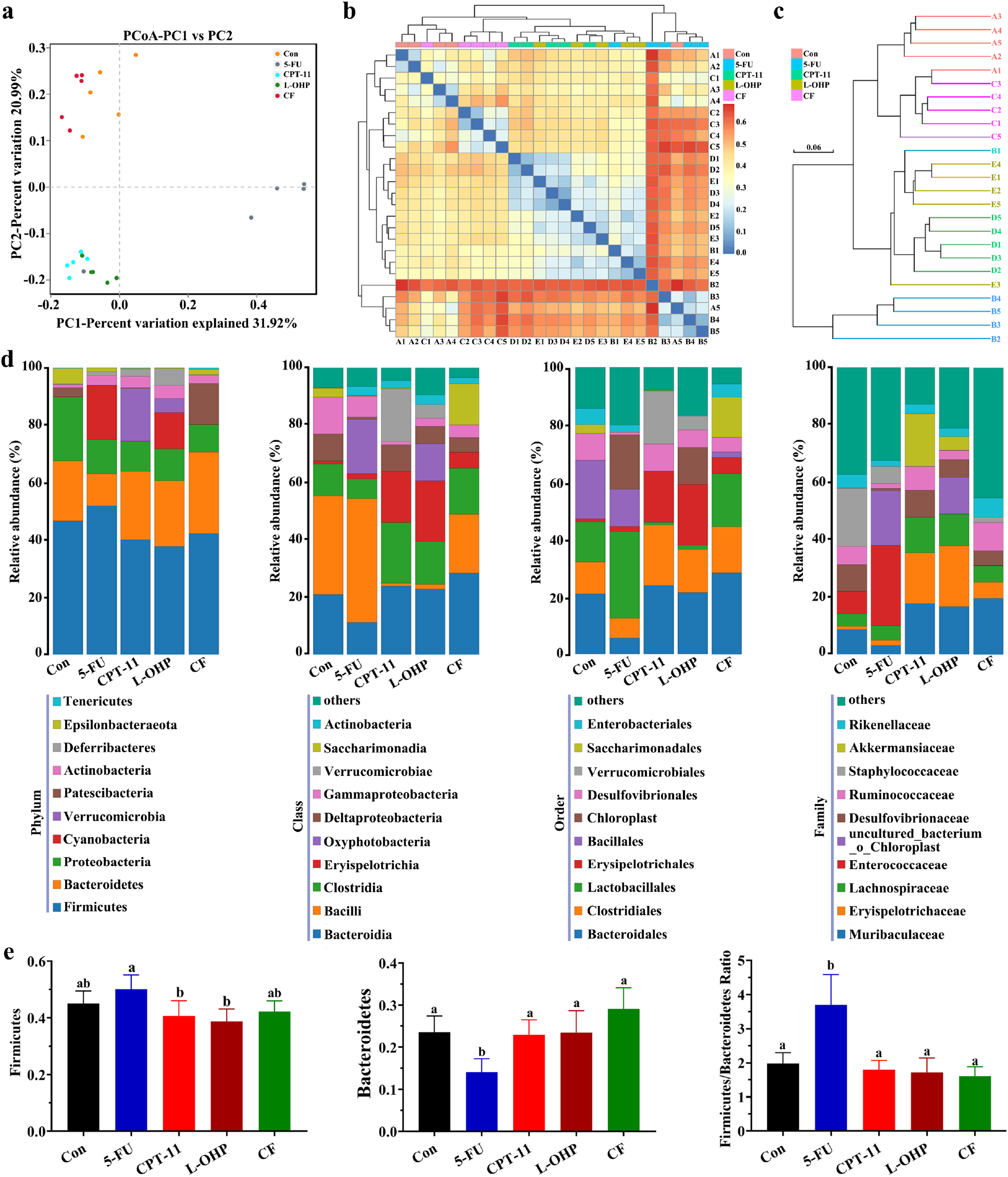
Effects of different chemotherapy drugs on the gut microbiota in mice. (a to c) Principal coordinate analysis (PCoA) (a), heatmap results of the weighted UniFrac distance (b) and arithmetic mean tree of the unweighted UniFrac distance (c) of microbial 16S rRNA sequences from the V3-V4 region in feces. (d) Average composition at the phylum, class, order, and family levels in the groups. (e) Graphs of gut microbiota composition was constructed using the levels of Firmicutes and Bacteroidetes as well as the Firmicutes to Bacteroidetes (F/B) ratio. One-way ANOVA followed by Tukey’s test was used to evaluate the statistical significance. The different letters represent significant differences between different groups (p < 0.05).

The results of the binary-based principal coordinate analysis (PCoA) showed distinct clustering of the microbiota composition in each group (Fig. 4a and S3). Compared with the control group, there was no significant deviation in the CF group, but not in the L-OHP, CPT-11 or 5-FU groups. The UniFrac-based weighted heatmap (Fig. 4b) and unweighted pair-group method with arithmetic mean tree (Fig. 4c) revealed that the microbial communities in the feces of the 5-FU, CPT-11 and L-OHP groups were significantly different from those in the control group. In addition, the samples of the 5-FU and CPT-11 groups were separated into two independent groups, while the samples of the L-OHP group were hybridized in the CPT-11 and 5-FU groups. These results suggested that the changes of gut microbiota in the 5-FU and CPT-11 groups were two different types, while the changes in the L-OHP group was somewhere between the 5-FU and CPT-11 groups.

To profile the specific changes in the gut microbiota, we analyzed the relative abundance of the predominant taxa (top 10) (Fig. 4d and e). At the phylum level, *Proteobacteria* significantly decreased and *Actinobacteria* significantly increased in 5-FU, CPT-11 and L-OHP groups (p < 0.05). *Patescibacteria* significantly decreased and *Cyanobacteria* significantly increased in 5-FU and L-OHP groups (p < 0.05). *Epsilonbacteraeota* decreased significantly, while *Verrucomicrobia* and *Deferribacteres* increased significantly in CPT-11 and L-OHP groups (p < 0.05, vs. control). The Firmicutes/Bacteroidetes (F/B) ratio was used as a common parameter to assess the gut microbiota disorder in many diseases (22). As shown in Fig. 4d to e, the F/B ratio in the 5-FU group significantly increased compared to the control group (p < 0.05), indicating that the components of the gut microbiota were harmful. However, no significant F/B ratio chang were found in L-OHP, CPT-11 and CF groups.

To identify bacterial taxonomic markers associated with intestinal injury and determine whether there were overlaps among the microbial change patterns under different chemotherapy drugs treatment, we constructed Fig. 5a, which was based on the results of line discriminant analysis (LDA) effect size (LEfSe) analysis and covered the significant changes of bacteria at all levels of phylum, class, order, family and genus as compared to controls (the name of bacteria corresponding to each number is shown in Table 1). From the Fig. 5a and table 1, we found that CF had little influence on the gut microbiota, only 8 bacteria significantly increased and 3 decreased. While, the L-OHP had the greatest influence on the gut microbiota, 31 bacteria significantly increased and 25 bacteria significantly decreased.

**FIG 5.**
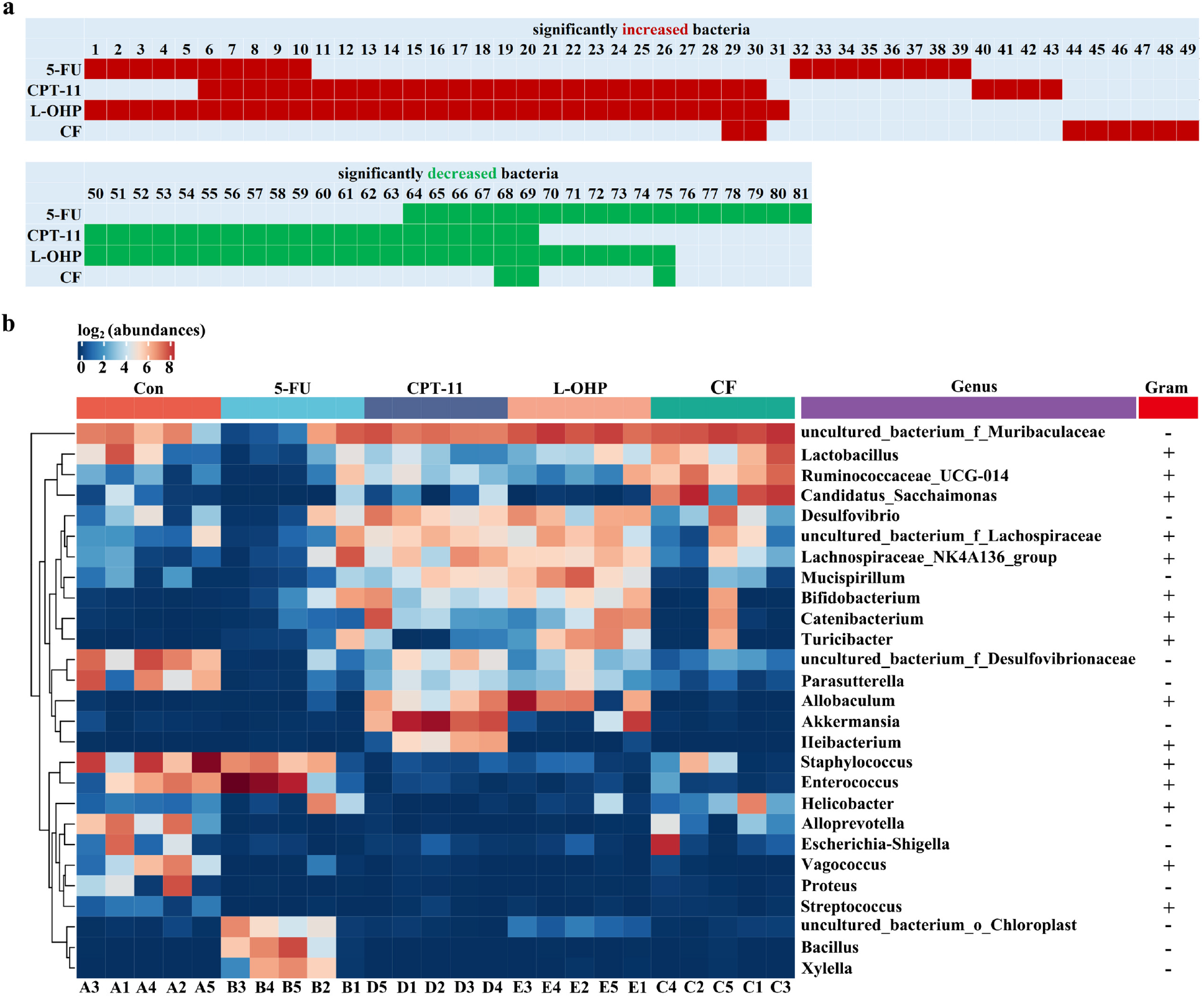
Effects of different chemotherapy drugs on microbiota taxonomic distributions in mice. (a) The significant changes of bacteria at all levels of phylum, class, order, family and genus as compared to controls. . (b) The heatmap of 16S rRNA gene sequencing analysis of feces at the genus level showed 27 key species with significant differences.

**Table 1.**

| NO. | Biological classification & Bacteria Name | NO. | Biological classification & Bacteria Name | NO. | Biological classification & Bacteria Name |
| --- | --- | --- | --- | --- | --- |
| **1** | p__Cyanobacteria | **28** | g__Lachnospiraceae_NK4A136_group | **55** | o__Enterobacteriales |
| **2** | c__Oxyphotobacteria | **29** | f__Muribaculaceae | **56** | o__Lactobacillales |
| **3** | o__Chloroplast | **30** | g__uncultured_bacterium_f__Muribaculaceae | **57** | f__Enterobacteriaceae |
| **4** | f__uncultured_bacterium_o__Chloroplast | **31** | g__Turicibacter | **58** | f__Enterococcaceae |
| **5** | g__uncultured_bacterium_o__Chloroplast | **32** | o__Sphingobacteriales | **59** | f__Helicobacteraceae |
| **6** | p__Actinobacteria | **33** | o__Xanthomonadales | **60** | f__Staphylococcaceae |
| **7** | c__Actinobacteria | **34** | c__Alphaproteobacteria | **61** | g__Enterococcus |
| **8** | o__Bifidobacteriales | **35** | f__Bacillaceae | **62** | g__Helicobacter |
| **9** | f__Bifidobacteriaceae | **36** | f__Sphingobacteriaceae | **63** | g__Staphylococcus |
| **10** | g__Bifidobacterium | **37** | f__Xanthomonadaceae | **64** | f__Prevotellaceae |
| **11** | p__Deferribacteres | **38** | g__Bacillus | **65** | g__Proteus |
| **12** | p__Verrucomicrobia | **39** | g__Xylella | **66** | g__Escherichia_Shigella |
| **13** | c__Deferribacteres | **40** | c__Clostridia | **67** | g__Alloprevotella |
| **14** | c__Erysipelotrichia | **41** | o__Clostridiales | **68** | g__Vagococcus |
| **15** | c__Verrucomicrobiae | **42** | f__Lachnospiraceae | **69** | g__Streptococcus |
| **16** | o__Deferribacterales | **43** | g__Ileibacterium | **70** | p__Patescibacteria |
| **17** | o__Erysipelotrichales | **44** | p__Patescibacteria | **71** | c__Saccharimonadia |
| **18** | o__Verrucomicrobiales | **45** | c__Saccharimonadia | **72** | o__Saccharimonadales |
| **19** | f__Akkermansiaceae | **46** | o__Saccharimonadales | **73** | f__Saccharimonadaceae |
| **20** | f__Deferribacteraceae | **47** | f__Saccharimonadaceae | **74** | g__Candidatus_Saccharimonas |
| 21 | f__Erysipelotrichaceae | 48 | g__Candidatus_Saccharimonas | 75 | g__uncultured_bacterium_f_Desulfovibrionaceae |
| 22 | g__Akkermansia | 49 | g__Ruminococcaceae_UCG_014 | 76 | c__Deltaproteobacteria |
| 23 | g__Allobaculum | 50 | p__Epsilonbacteraeota | 77 | o__Desulfovibrionales |
| 24 | g__Catenibacterium | 51 | c__Bacilli | 78 | f__Desulfovibrionaceae |
| 25 | g__Mucispirillum | 52 | c__Campylobacteria | 79 | f__Lactobacillaceae |
| 26 | g__Desulfovibrio | 53 | o__Bacillales | 80 | g__Lactobacillus |
| 27 | g__uncultured_bacterium_f_Lachnospiraceae | 54 | o__Campylobacteriales | 81 | g__Parasutterella |

The microbial change pattern of L-OHP group intersected with CPT-11 and 5-FU groups. 29 bacteria were significantly increased under CPT-11 treatment, and 25 of these (86%) were also increased in the L-OHP group. In addition, 19 bacteria significantly decreased under CPT-11 treatment and all of them (100%) were also decreased in the L-OHP group. Interestingly, 18 bacteria were significantly increased under 5-FU treatment, and 10 of these (55.6%) were also changed in the L-OHP group. 17 bacteria significantly decreased under 5-FU treatment, and 11 (64.7%) of them were also decreased in the L-OHP group (Fig. 5a). In addition, as show in Fig. 5a the microbial change pattern of 5-FU and CPT-11 were almost two different types.

A genus level heatmap with the relative abundance of 27 genera was constructed to show the change in the gut microbiota caused by 5-FU, CPT-11, L-OHP, or CF (Fig. 5b). The heatmap of the relative abundance of microorganisms altered by the four chemotherapy drugs showed a difference in gut bacterial composition compared to that of the control group at the genus level.

### The molecular mechanism of gut microbiota disorganization leads to the aggravation of intestinal injury

We further used phylogenetic investigation of communities by reconstruction of unobserved states, V1.0 (PICRUSt) to predict the metabolic functions according to the gut microbiota changes under four chemotherapy drugs treatment. The relative level of “bacterial chemotaxis”, “antigen processin and presentation”, and “NOD like signaling pathway” were significantly higher in the 5-FU, CPT-11 and L-OHP groups than in the control and CF groups (Fig. 6a). Activating the NOD like pathway promotes the secretion of inflammatory cytokines, and induces intestinal injury (23). Therefore, we hypothesized that the intestinal injury aggravation induced by the gut microbiota disorder under treatment with different chemotherapy drugs is associated with the NOD like signaling pathway.

**FIG 6.**
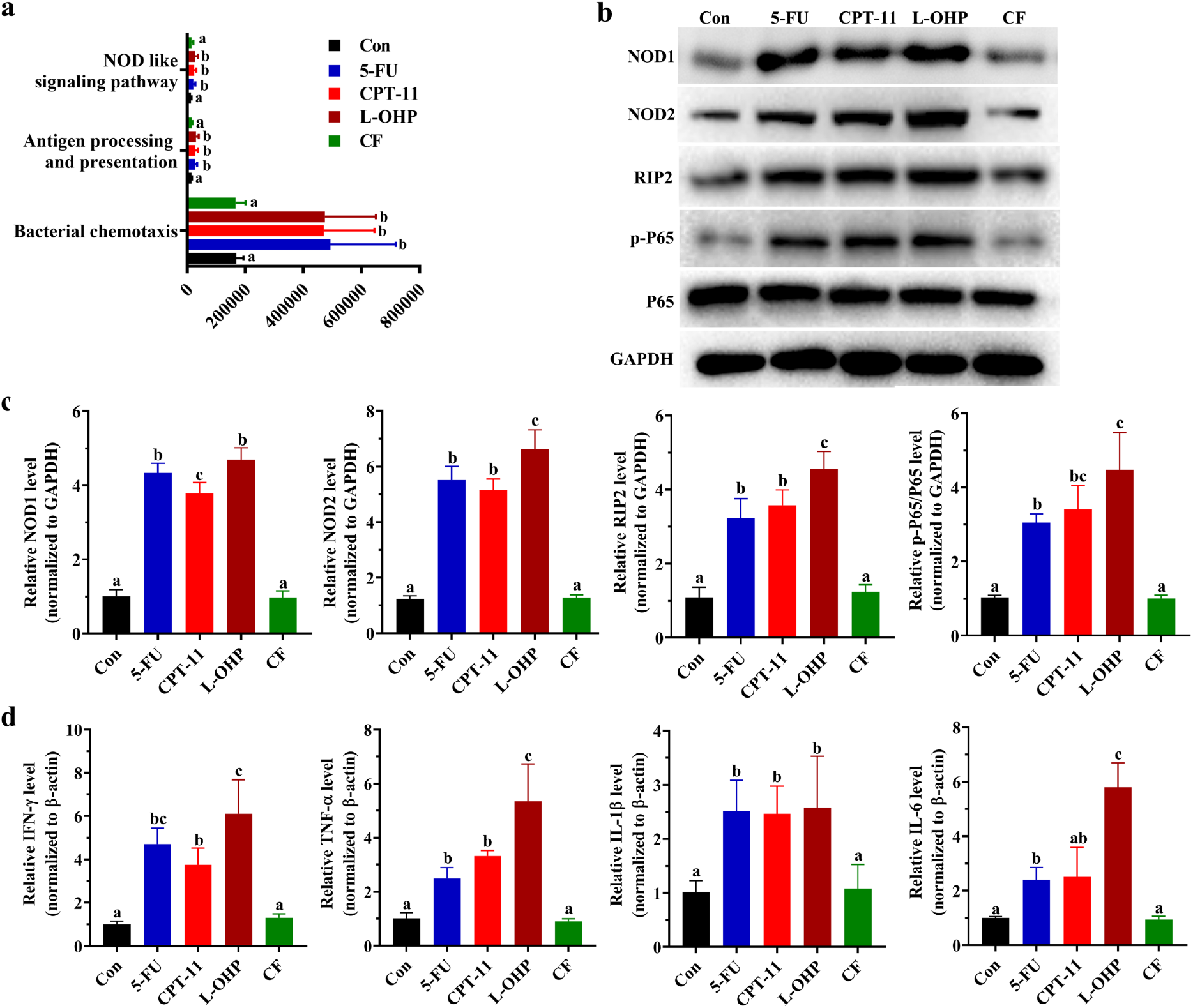
Effects of different chemotherapy drugs on the prediction of potential metabolic functions of the gut microbiota in mice and verify it. (a) Microbial gene functions in the mice of different groups as indicated using PICRUSt bioinformatics software package. The different letters represent significant differences between different groups (p < 0.05). (b to d) The effect of different chemotherapy drugs on NOD1/2/RIP2/NF-κB pathway proteins (b and c) andinflammatory cytokines (d). One-way ANOVA followed by Tukey’s test was used to evaluate the statistical significance. The different letters represent significant differences between different groups (p < 0.05).

To test this hypothesis, we detected NOD1 and NOD2 gene expression at the protein levels in jejunum tissues from different groups and found that NOD1/2 levels increased in 5-FU, CPT-11 and L-OHP groups (p < 0.05) (Fig. 6b to c). The activation of NOD1/2 can activate the NODs/RIP2/NF-κB (p65) signaling pathway to motivate inflammation(24). We found that under 5-FU, CPT-11 and L-OHP treatment, the RIP2 and p-P65/P65 at the protein level were up-regulated (Fig. 6b and c). Furthermore, we detected the inflammatory cytokines in jejunum tissues by Q-PCR. We found that 5-FU, CPT-11 and L-OHP administration significantly enhanced the secretion of inflammatory cytokines in intestines, including IFN-γ, TNF-α, IL-1β, and IL-6 (p < 0.05) compared to the control group (Fig. 6d).

### Correlation analysis of biological indicators of the gut microbiota

Finally, according to the Pearson correction coefficient between 27 genera and 10 parameters (IFN-γ, IL-1β, Il-6, TNF-α, ZO-1, occludin, NOD1, NOD2, RIP2 and p-P65/P65), we created a correction matrix. It was found that 70.4% (19/27) of the 27 genera influenced by chemotherapy drugs were negatively or positively related to one or more parameters (Fig. 7). These data suggest that the gut microbiota plays an important role in chemotherapy drugs-induced intestinal injury and is significantly related to the secretion of inflammatory cytokines, the integrity of the intestinal mucosa, and the NODs/RIP2/NF-κB pathway.

**FIG 7.**
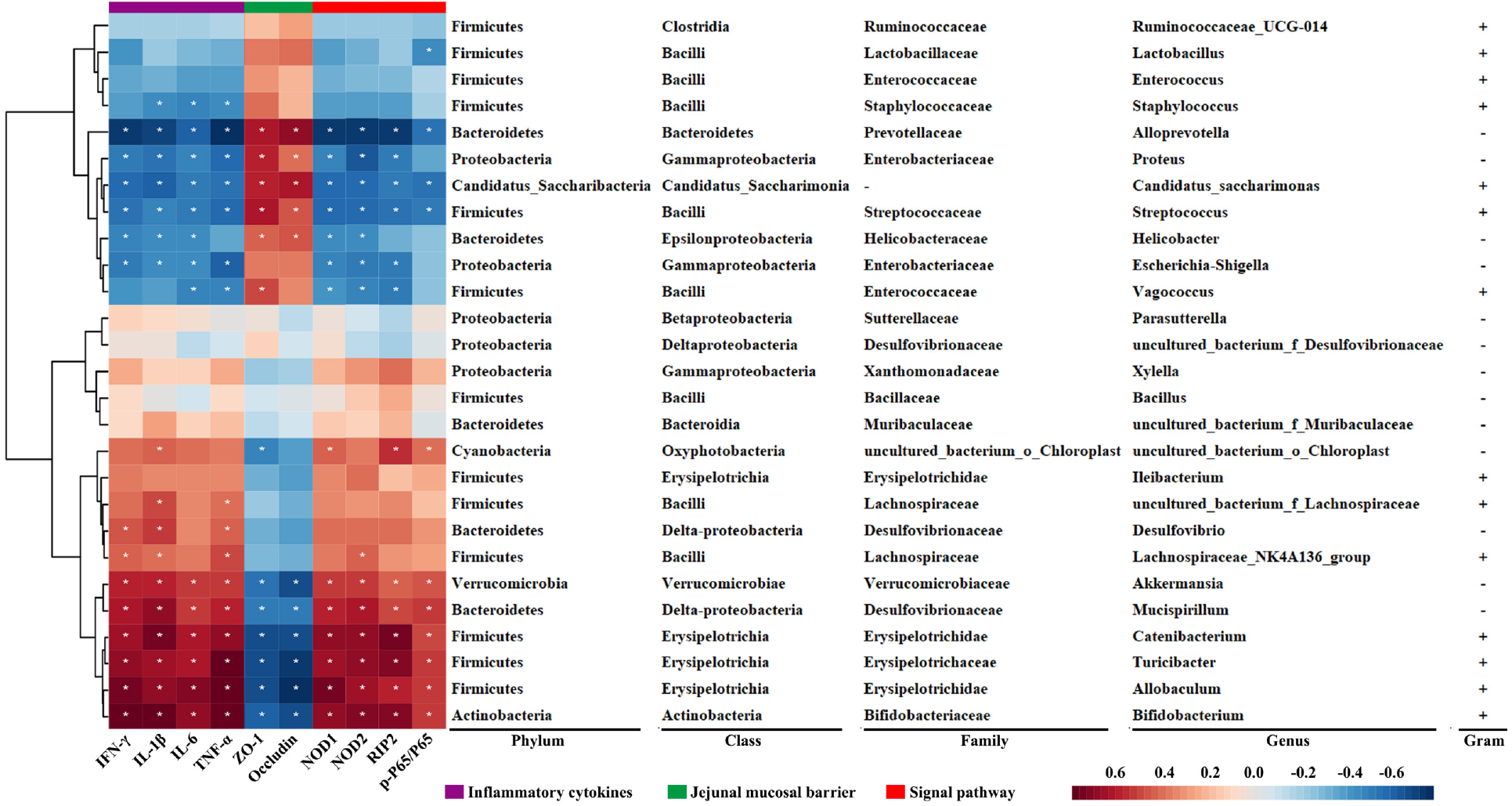
Correlation analysis of biological indicators related to the gut microbiota. The heatmap shows the value of the correlation coefficient between the gut microbiota and biological markers. Correlation coefficient R > 0.25 indicates a moderate correlation and R^2^ > 0.36 indicates a strong correlation. * and ** indicate the correlation significance (p < 0.05 and p < 0.01, respectively). Red represents a positive correlation and blue represents a negative correlation. The darker the color, the stronger the correlation. Each row shows bacterial taxa information (phylum, class, family, and genus).

## DISCUSSION

Chemotherapy drugs not only influence tumor cells but also normal cells. The use of chemotherapy drugs is usually accompanied by mucosal lesions because of the overproduction of proinflammatory cytokines and apoptosis of intestinal epithelial cells (25). The destruction of the intestines induced by chemotherapy drugs leads to a variety of side effects, such as abdominal pain, diarrhea, fecal blood, and even death in severe cases (9, 10). Therefore, clarifying the level of different chemotherapy drugs-induced intestinal injury as well as the main mechanism can help to develop targeted detoxification therapy and increase the clinical application of chemotherapy, which is meaningful for the treatment of colon cancer.

Although previous studies have shown that chemotherapy drugs, such as 5-FU, CPT-11 and L-OHP, can induce intestinal injury (26), none have compared the differences between the intestinal injury degree and gut microbiota disorder pattern among them, which form the FOLFIRI and FOLFOX for colorectal cancer chemotherapy regimens (27). Our study excluded the effects of different operators, seasonal climates, batches of mice, and mice from different environments, which allowed us to obtain accurate results for 5-FU-, CPT-11-, L-OHP-, and CF-induced intestinal injury and gut microbiota disorder in BALB/c mice. Comparing these results will allow the development of new ideas and a scientific theory for reducing the intestinal side effects of colorectal cancer chemotherapy.

5-FU, an S-phase-specific anticancer drug, causes intestinal injury in 50-80% of patients (28). L-OHP, a third-generation platinum antitumor compound, can for minter- or intra-strand crosslinks in DNA to prevent DNA replication and transcription, resulting in apoptosis. The common adverse reaction of L-OHP is colon toxicity, and the diarrhea dose of L-OHP overlaps with its lethal dose (5). Chemotherapy with CPT-11 also induces intestinal mucositis with increased expression of proinflammatory cytokines regulated by NODs (29). CF alone does not inhibit tumor growth, but in combination with 5-FU, it increases the antitumor effect of 5-FU (30).

In this study, we found that mice under 5-FU or CPT-11 treatment displayed moderate weight loss, diarrhea, and fecal blood symptoms. L-OHP treatment caused most severe damage than 5-FU and CPT-11 (Fig. 1). 5-FU, CPT-11 and L-OHP could induce colon shortening, significantly induce colon and jejunum injury, and the most severe intestinal injury was caused by L-OHP. In addition, only L-OHP could cause significant histopathological injury to the liver and spleen. Therefore, through morphological observations, we found that L-OHP had the most severe side effects and caused the most serious intestinal injury among the main colorectal cancer chemotherapy drugs (Fig. 2).

Caspase-3 is a key executor of apoptosis, which can be cleaved to its activated form, and induces cell apoptosis. The movement of Bax from the cytosol to the mitochondria induces the release of cytochrome C and activation of caspase-3, which can be inhibited by Bcl-2 (31). Administration of 5-FU, CPT-11 and L-OHP increased the protein levels of Bax and cleaved caspase-3, but decreased the protein level of Bcl-2. However, the protein levels of the above-mentioned molecules in the CF group were not significantly different from those in the control group (Fig. 3). As the first line of immune defense, the intestinal epithelial barrier is crucial for protecting the host against invasive pathogenic bacteria. Once the integrity of the intestinal barrier is lost, bacteria and toxic substances can penetrate through the intestinal wall and trigger the aforementioned feedback cycle (32). We found that 5-FU, CPT-11 and L-OHP significantly decreased the expression of ZO-1 and occluding (Fig. 3), markers of mucosal barrier integrity in the jejunum (33).

We analyzed the composition of the intestinal microbiota in the feces. PCoA results revealed that despite access to the same food, the microbial communities under 5-FU, CPT-11 or L-OHP treatment varied considerably. The beta-diversity heatmap and ClusterTree analyses results were in accordance with those of PCoA, and showed that the microbial communities under L-OHP treatment were more similar to CPT-11 than 5-FU (Fig. 4). The F/B ratio was used as a common parameter to assess the degree of intestinal microbiota disturbance (34). In this study, at the phylum level, the F/B ratio was increased in the 5-FU group (P<0.05, vs. control), while no significant difference was observed in CPT-11, L-OHP and CF groups (P>0.05, vs. control). The change in the F/B ratio confirmed that the type of gut microbiota change induced by CPT-11 and L-OHP was different from 5-FU. The relative abundance of microbiota at the class, order, and family levels was also consistent with this phenomenon (Fig. 4).

To provide guidelines for the targeted treatment of chemotherapy -induced intestinal injury, we carefully compared the change pattern of 5-FU, CPT-11, L-OHP and CF groups. Using LEfSe to identify microbial biomarkers. 86% (25/29) of the increased bacteria and 100% (19/19) of the decreased bacteria under CPT-11 treatment appeared in the change pattern under L-OHP treatment. However, only 17.2% (5/29) of the increased bacteria and 26.3% (5/19) of the decreased bacteria under CPT-11 treatment appeared in the 5-FU’s microbial change pattern. In the microbial change pattern under 5-FU treatment, 55.6% (10/18) of the increased bacteria and 64.7% (11/17) of the decreased bacteria appeared in the L-OHP group (Fig. 5). In combination with the intestinal injury test, L-OHP was found to cause most severe injury than either 5-FU or CPT-11(Fig. 1 and 2). So, we hypothesized that L-OHP caused the most severe intestinal injury because of the most extensive microbial change pattern, and the pattern is overlap with CPT-11 and 5-FU.

Based on these results, we further compared the influence of 5-FU, CPT-11, L-OHP, and CF on genus-level taxonomic distributions of the microbial communities in feces (Fig. 5). After 5-FU treatment, the abundance of *Lactobacillus, Parasutterella*, *Alloprevotella*, *Escherichia-Shigella*, *Vagococcus*, *Proteus*, *uncultured bacterium f Desulfovibrionaceae*, and *Candidatus Saccharimonas* sp. was significantly reduced, while the abundance of *Bifidobacterium*, *uncultured bacterium o Chloroplast*, *Bacillus*, and *Xylella* sp. was significantly increased. Gram-positive bacteria *Lactobacillus* and *Parasutterella* sp. are beneficial bacteria, and *Parasutterella* sp. are related to short-chain fatty acid (SCFA) production (35). The Gram-negative bacteria *Alloprevotella* sp. are also SCFA-producing bacteria (36). *Escherichia-Shigella* sp. are associated with inflammation in both the acute necrotizing pancreatitis (ANP) model (37) and intestinal inflammation model (38). The abundance of intestinal *Vagococcus* sp. in healthy individuals is significantly higher than that in patients with cancer (39). *Proteus* sp. are usually considered commensals in the gut and are recognized clinically as putative gastrointestinal pathogens (40). The abundance of *Candidatus Saccharimonas* decreased significantly in the ANP model (37, 41), but increased significantly in the colitis-associated carcinogenesis model (42). *Bifidobacterium* sp. are probiotics that protect the neonatal intestine against necrotizing enterocolitis (43). Some species of *Bacillus* are human pathogens, such as *Bacillus cereus*, *Bacillusanthracis*, and *Bacillus mycoides* (44). *Xylella* are common pathogens in plants (45), but their abundance significantly increased with the intervention of 5-FU.

After CPT-11 treatment, the abundance of *Vagococcus*, *Proteus*, *Escherichia-Shigella*, *Alloprevotella*, *Enterococcus*, *Helicobacter*, and *Staphylococcus* sp. significantly decreased, while that of *Akkermansia*, *Allobaculum*, *Ileibacterium*, *Catenibacterium*, *Bifidobacterium*, *Desulfovibrio*, *Lachnospiraceae NK4A136 group, Mucispirillum, uncultured bacteriumf Lachnospiraceae*, and *uncultured bacteriumf Muribaculaceae* sp. significantly increased. *Enterococcus* sp. are a common pathogen of prostatitis (46), but they are not highly virulent. *Helicobacter* sp. are potent drivers of colonic T cells and are positively correlated with body weight (36). The majority of *Staphylococcus* sp. are non-pathogenic bacteria and several can cause disease (47). *Akkermansia* sp. are considered a probiotic in most studies, but the presence of excessive *Akkermansia* sp. can cause intestinal injury (48). The excessive abundance of *Catenibacterium, Bifidobacterium*, and *Allobaculum* sp. had a significantly positive relationship with serum inflammatory factors and may aggravate the severity of colitis (49, 50), and *Ileibacterium* were newly identified bacteria of the genus *Allobaculum* (51). *Desulfovibrio* and *Lachnospiraceae NK4A136 group* sp. are harmful bacteria associated with a high-fat diet and chronic inflammation (52, 53). *Mucispirillum* sp. inhabit the mucus layer over colonocytes and are associated with the early disruption of the colonic surface mucus layer (54).

Significantly changed gut microbiota under L-OHP treatment almost appeared in the microbial change pattern of CPT-11 or 5-FU group, except *Turicibacter* sp., which aggravates the severity of colitis (49). The abundance of *Vagococcus*, *Proteus*, *Escherichia-Shigella*, and *Alloprevotella* sp. decreased in the L-OHP, CPT-11, and 5-FU groups. *Enterococcus*, *Helicobacter*, and *Staphylococcus* sp. levels decreased in the L-OHP and CPT-11 groups. *Candidatus*-*Saccharimonas* and *uncultured bacterium f Desulfovibrionaceae* sp. decreased in the L-OHP and 5-FU groups. *Bifidobacterium* increased in the L-OHP, CPT-11, and 5-FU groups. *Desulfovibrio*, *Lachnospiraceae_NK4A136_group*, *Mucispirillum*, *Akkermansia*, *Allobaculum*, *Catenibacterium*, *uncultured bacteriumf Lachnospiraceae*, and *uncultured bacteriumf Muribaculaceae* sp. increased in the L-OHP and CPT-11 groups. *Uncultured bacterium o Chloroplast* sp. increased in the L-OHP and 5-FU groups.

Differences in the functional genes of the gut microbiota in metabolic pathways between different chemotherapy drugs groups were predicted by PICRUSt. The influence of the gut microbiota on the host cellular processes, organismal systems, and environmental information processing may lead to the occurrence of diseases. We found that the influence of 5-FU, CPT-11 and L-OHP on the gut microbiota was found to be related to “Bacterial chemotaxis”, “Antigen processing and presentation”, and “NOD like signaling pathway” (Fig. 6). “Bacterial chemotaxis” is the phenomenon whereby bacteria direct their movements according to certain chemical stimulants in their environment (55), and the “Bacterial chemotaxis” helps harmful bacteria to colonize the intestinal mucosa and promote intestinal injury (56). NODs mediate the innate immune system and regulate the expression of proinflammatory cytokines (57). “NOD like signaling pathway” activated by intestinal mucosal injury may induce immune responses and aggravate the severity of the injury (58). NODs/RIP2/NF-κB signaling patheway is closely associated with oxidative stress and inflammatory responses (59, 60), and NF-κB can alter the expression of inflammatory cytokines, such as IFN-γ, TNF-α, IL-1β and IL-6, to cause inflammatory injury in the intestinal barrier (61). These results suggest that chemotherapy drugs may further activate the expression of proinflammatory cytokines to promote intestinal injury by activating the NODs/RIP2/NF-κB signaling pathway. The protein expression of NOD1, NOD2, RIP2, and p-NF-κB/ NF-κB, and the secretion level of proinflammatory cytokines in intestines confirmed this hypothesis (Fig. 6). Furthermore, based on the correlation analysis of intestinal injury biological indicators and the gut microbiota, we found that 19 of the 27 genera (70.3%) were associated with at least 1 parameter, and 13 of the 27 genera (48%) were associated with aleast 6 parameters (Fig. 7).

Taken together, chemotherapy drugs for colorectal cancer, 5-FU, CPT-11, L-OHP and CF, have different toxicities to intestinal injury and gut microbiota, and L-OHP induced the most severely damage. The microbial change pattern under L-OHP treatment had significant overlap with the change in 5-FU and CPT-11 groups, but the microbial change pattern didn’t overlap between 5-FU and CPT-11. The NODs/RIP2/NF-κB signaling pathway was also maximum activated under L-OHP treatment. 13 Genera of particular interest were *Bifidobacterium*, *Akkermansia*, *Allobaculum*, *Catenibacterium*, *Mucispirillum*, *Turicibacter*, *Helicobacter*, *Proteus*, *Escherichia Shigella*, *Alloprevotealla*, *Vagococcus*, *Streptococcus* and *Candidatus Saccharimonas* sp. showed different abundances in the four drugs groups and were most likely correlated with the mechanism of intestinal injury. The above results may explain why different chemotherapy drugs induce different levels of intestinal injury (Fig. 8), and suggest the use of bacteriostatic agents should be targeted to different colorectal cancer chemotherapy drugs, and provide new ideas and new research directions for the targeted treatment of intestinal injury caused by colorectal cancer chemotherapy,.

**FIG 8.**
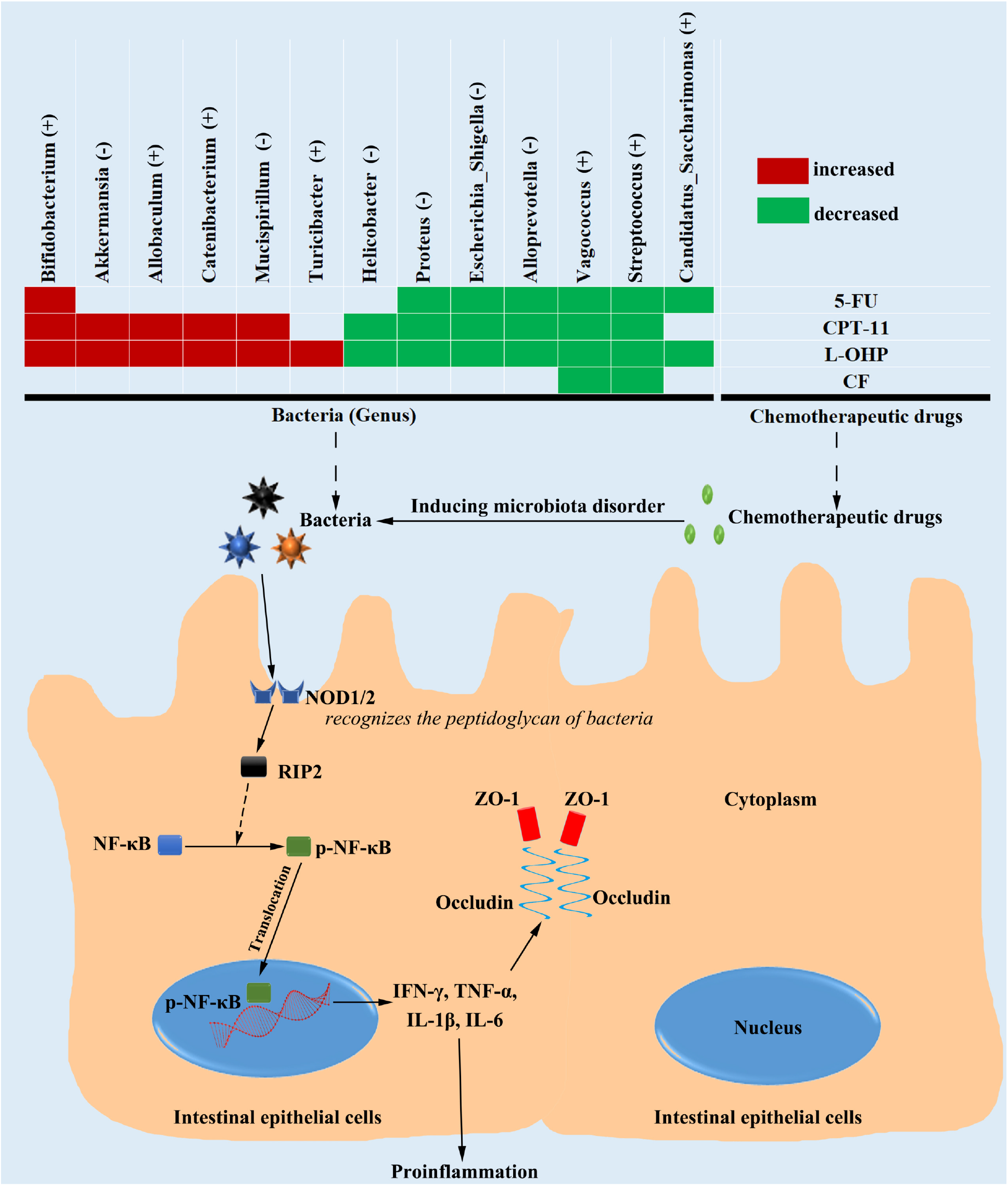
Mechanisms of different degrees of intestinal injury induced by chemotherapy drugs (5-FU, CPT-11, L-OHP, CF) for colorectal cancer.

## MATERIALS AND METHODS

### Cell culture

CT-26 cells were purchased from the Cell Bank of the Chinese Academy of Sciences (Shanghai, China). Cells were cultured in Roswell Park Memorial Institute 1640 medium (C11875500BT, Life Technologies, Carlsbad, CA, USA) containing 10% (v/v) fetal bovine serum (10099141, Life Technologies) and 1% penicillin-streptomycin (SV30010, Life Technologies) at 37°C in a 5% CO_2_ humidified incubator.

### Construction of an intestinal injury model induced by chemotherapy drugs in tumor-bearing mice

Four-week-old male BALB/c mice weighing 20-24 g were obtained from the SLAC Laboratory Animal Technology Co., Ltd (Shanghai, China). The mice were maintained under specific pathogen-free conditions (12 h light/dark cycle; 23-25°C; 45-55% relative humidity) and were given ad libitum access to autoclaved food and water. After one week of pre-experimental adaptation, all mice were subcutaneously injected with 4×10^6^ CT-26 cells in the right armpit. Tumor-bearing mice were randomly divided into five experimental groups (n=8) as follows: control, 5-FU, CPT-11, L-OHP, and CF groups. According to previous studies (18) and our exploration of pre-project modeling conditions, the chemotherapy clinical equivalent dose was used as the intervention dose to induce chemotherapy-induced intestinal injury. The operation is shown in Fig. 1a. The mice in different groups were respectively intraperitoneally administered 200 μL of saline (control), 5-FU (100 mg/kg, R050245; RHAWN, Shanghai, China), CPT-11 (85 mg/kg, B21490; Shanghai Yuanye Bio-Technology Co., Ltd, Shanghai, China), L-OHP (20 mg/kg, R002057; RHAWN, Shanghai, China), and CF (100 mg/kg, R054174; RHAWN, Shanghai, China) for five consecutive days.

Body weight, diarrhea index, FOB score, and number of mortalities were recorded daily. The severity of body weight loss, diarrhea index, and FOB was assessed according to Table S1. FOB was detected using the Baso OB Kit (BA2020B, Zhuhai, China). Then, the DAI (FOB + diarrhea index + body weight loss index) was calculated. All mice were anesthetized with sodium pentobarbital and euthanized after a 16 h fast on Day 6. Next, the jejunum and colon were cut into 1 cm-length sections, and one tissue was fixed in 4% paraformaldehyde for histological or immunohistochemical evaluation. The other jejunal tissues for RNA-seq analysis, western blotting, and Q-PCR were frozen at -80°C. Sera were collected for the detection of proinflammatory cytokines.

All animal-related protocols of this study conformed to the rules of the Animal Experimental Ethics Committee of Fujian University of Traditional Chinese Medicine and were performed in accordance with the National Institutes of Health Guidelines for the care and use of laboratory animals.

### Histological analysis of the intestinal injury

The jejunal and colonic tissues were collected, fixed in 4% paraformaldehyde, and embedded in paraffin. These were then cut into 4 μM-thick sections to prepare slides and were then stained with hematoxylin and eosin (HE) (Solarbio Science, Beijing, China). Pathological changes were observed under a light microscope (Leica DM4000B; Leica Co., Germany). The villus height and crypt depth of the jejunum and colon were measured in at least 10 clear longitudinal sections.

### Enzyme-linked immunosorbent assay (ELISA)

The levels of IFN-γ, TNF-α, IL-1β, IL-6, and LPS in the sera of each mouse were detected according to the ELISA kit protocol (Meimian, Jiangsu, China).

### Immunohistochemical evaluation

PCNA expression was detected using immunohistochemistry (IHC) analysis. The intestinal slides were deparaffinized and rehydrated. Endogenous peroxidase was blocked with 3% H_2_O_2_. Sections were incubated overnight at 4°C with the primary antibodies; anti-PCNA antibody (1:200, Sanying biotechnology, Hubei, China) and the secondary antibodies (1:200, Sanying biotechnology, Hubei, China), and then stained with DAB and hematoxylin. Sections were observed under a light microscope (Leica). The integrated optical density of the sections positively-stained for the proteins was measured using ImageJ software, version 1.8.0.

### Assessment of apoptosis (TUNEL staining)

For the assessment of apoptosis, slides of jejunum sections were processed for the TUNEL assay (MK1025; Boster, Hubei, China). Images were taken using a microscope (Leica) at both 200× and 400× magnification.

### Western blot analysis

Jejunum tissues were harvested and total protein was extracted using radioimmunoprecipitation assay lysis buffer containing proteinase inhibitor (Beyotime, Haimen, China). Total protein content was determined using the BCA protein assay (Beyotime). After quantification, protein samples were separated by SDS-PAGE, transferred to PVDF membranes, and incubated with one primary antibody (antibodies are detailed in Supplementary Table S2) and HRP-conjugated secondary antibodies. Chemiluminescent detection was conducted using the ChemiDoc XRS+ imaging system (Bio-Rad Laboratories, Inc., Hercules, CA, USA) according to the manufacturer’s instructions. ImageJ software was used to determine protein expression levels.

### Q-PCR

Total RNA was extracted from jejunum tissues using RNA isolation reagent (TaKaRa Biotechnology, Dalian, Liaoning, China) according to the manufacturer’s instructions for animal tissue. Subsequently, the total RNA was reverse transcribed to cDNA using the Prime Script_TM_ RT reagent kit (TaKaRa Biotechnology). qPCR was performed using the SYBR Premix Ex Taq II Kit (TaKaRa Biotechnology, Dalian, Liaoning, China) on ABI 7500 Sequence Detection System software version 1.2.3 (Applied Biosystems, CA, USA). The thermocycling conditions were as follows: 50°C for 2 min, 95°C for 10 min, and 40 cycles of 95°C for 15 s and 60°C for 1 min. β-Actin was used as an endogenous control. The 2^-ΔΔCT^ method was used to calculate the TLR4 mRNA levels.

### Gut microbiota analysis

**(i) DNA extraction** Feces were collected from five mice per group and immediately stored at -80°C. DNA from feces was extracted using MN NucleoSpin 96 Soil (MN, 740787) according to the manufacturer’s instructions. **(ii) Sequence analysis.** Hypervariable region V3rer instructions. The DNA concentration was determined using a NanoDrop 2000 Spectrophotometer (Thermo Scientific, 341F: CCTAYGGGRBGCASCAG; 806R: GGACTACNNGGGTATCTAAT). Library construction and amplicon sequencing were performed using Genomics BioScience. A paired-end library (insert size of 460 bp for each sample) was constructed using the TruSeq Nano DNA Library Prep kit (Illumina), and high-throughput sequencing was performed on an Illumina HiSeq 2500 sequencer with a HiSeq Rapid v2 Reagent Kit (Illumina). The sequences of 2 × 250 bp paired-end reads were produced from the sequencer following the manufacturer’s instructions. The pair-reads were merged into amplicon sequences using FLASH v1.2.7, and these amplicon sequences were checked for the existence of the primers, duplicates were removed, and short sequences and chimeric reads were filtered out to generate effective reads. Effective reads were analyzed to generate operational taxonomic units (OTUs). Further 16S rDNA analysis (OTU picking and taxonomic assignment) and data visualization were conducted using Quantitative Insights Into Microbial Ecology (QIIME) version 1.8.0, with the Silva 16S rRNA Taxonomy Database (Release132). Taxonomy (i.e., phyla and OTUs) was analyzed using one-way analysis of variance (ANOVA). **(iii) Microbial analyses.** To clarify the diversity of intestinal microbiota composition among all experimental groups (control, 5-FU, CPT-11, L-OHP, and CF groups), the PCoA of unweighted UniFrac distance matrix are displayed as β-diversity. Weighted UniFrac distance matrices were computed to detect global variations in the composition of microbial communities at the phylum and genus levels using one-way ANOVA. Microbiota-based biomarker discoveries were performed using LEfSe. The relative abundance of each biomarker taxon across all samples is shown with straight and dotted lines that plot the means and medians, respectively, in each group. Color codes indicate the groupsand letters indicate the taxa that contribute to the uniqueness of the corresponding groups at LDA > 4.0. **(iv) Functional capacity of the microbiota.** PICRUSt was used to predict the 16S rRNA-based high-throughput sequencing data for functional features from the phylogenetic information with an estimated accuracy of 0.8. The cluster of orthologous group database was obtained through the Green-gene ID corresponding to each OTU in the EGGNOG database (evolutionary genealogy of genes: non-supervised orthologous groups). The predicted functional composition profiles were then collapsed into class 3 of the Kyoto Encyclopedia of Genes and Genomes (KEGG) database pathways. The correlations between 27 genera and the intestinal mucosal barrier (ZO-1 and occludin), inflammatory cytokines (IFN-γ, TNF-α, IL-1β, and IL-6), and PI3K/AKT/NF-κB signaling pathway-related genes in mice treated with different chemotherapy drugs were determined by RDA/CCA analysis and the related heatmap.

### Statistical analysis

Statistical analysis was performed using SPSS software (version 23.0; SPSS, Inc., Chicago, IL, USA). Data are presented as mean ± SD. Differences were analyzed using one-way ANOVA followed byan unpaired *t*-test. Pearson’s correlation was used to determine the relationships between the parameters. p < 0.05 was considered statistically significant. There was no statistically significant (p > 0.05) between groups with the same letter on the bar and there was statistically significant (p < 0.05) with the different letter.

## SUPPLEMENTAL MATERIAL

**Supplemental material available online only.**

**FIGS1,TIF file,3MB.**

**FIGS2,TIF file,12MB.**

**FIGS3,TIF file,342KB.**

## ACKNOWLEDGMENTS

This work was supported by the Natural Science Foundation of Fujian Province, China (2021J01907, 2021J01941), and the Developmental Fund of Chen Keji Integrative Medicine (No. CKJ2021006 and CKJ2021008).

---

**FIG S1** Photomicrographs of H&E-stained liver, spleen, renal medulla and renal cortex sections.

**FIG S2** Photomicrographs of H&E-stained colon sections.

**FIG S3** PCoA of different groups.

**Table S1.**

| **Score** | **Body Weight Loss** | **Diarrhea Index** | **Time Cost for detection of Fecal blood** |
| --- | --- | --- | --- |
| 0 | $\leq 1\%$ | normal stool | $\geq 2$ min |
| 1 | 1-5% | slightly wet and soft stool | 1-2 min |
| 2 | 6-10% | soft cylinder and easily spreadable | 30 s-1 min |
| 3 | 11-15% | wet and unformed stool | 10 s-30 s |
| 4 | $\geq 15\%$ | non-cylindrical or runny | $\leq 10$ s or gross visibility of blood |

**Table S2:** Antibodies for western blot.

| **Antibody** | **Cat No.** | **WB** | **Company** |
| --- | --- | --- | --- |
| Zo-1 | 21773-1-AP | 1/2000 | Wuhan Sanying bio-technology co., Ltd., China |
| Occludin | 27260-1-AP | 1/2000 | Wuhan Sanying bio-technology co., Ltd., China |
| NOD1 | DF6378 | 1/1000 | Cell Signaling Technology , USA |
| NOD2 | DF12125 | 1/1000 | Cell Signaling Technology , USA |
| RIP2 | 15366-1-AP | 1/1000 | Wuhan Sanying bio-technology co., Ltd. , China |
| p-AKT | mAb #4060 | 1/1000 | Cell Signaling Technology, USA |
| AKT | mAb #4691 | 1/1000 | Cell Signaling Technology, USA |
| p-p65 | mAb #3033 | 1/1000 | Cell Signaling Technology, USA |
| p65 | mAb #8242 | 1/1000 | Cell Signaling Technology, USA |
| GAPDH | 60004-1-Ig | 1/5000 | Wuhan Sanying bio-technology co., Ltd., China |
| Mouse IgG [HpL] | SA00001-1 | 1/10000 | Wuhan Sanying bio-technology co., Ltd., China |
| Rabbit IgG [HpL] | SA00001-2 | 1/10000 | Wuhan Sanying bio-technology co., Ltd., China |

